# Neural Systems Underlying the Implementation of Working Memory Removal Operations

**DOI:** 10.1101/2023.02.14.519204

**Authors:** Jacob DeRosa, Hyojeong Kim, Jarrod Lewis-Peacock, Marie T. Banich

## Abstract

Recently multi-voxel pattern analysis has verified the removal of information from working memory (WM) via three distinct operations *replacement, suppression*, or *clearing* compared to information being *maintained* (Kim et al., 2020). Univariate analyses and classifier importance maps indicate that some brain regions commonly contribute to these operations. This study aimed to use multivariate approaches to determine whether, within these commonly activated brain regions, each of these operations is being represented in a similar or distinct manner. To do so, we used *Leiden community detection* to identify brain networks that are characterized by similar multi-voxel patterns of activity with regard to these WM operations. Four networks were identified. The Visual Network shows similar multi-voxel patterns for *maintain* and *replace*, which are highly dissimilar from *suppress* and *clear*, suggesting this network differentiates whether an item is held in WM or not. The Somatomotor Network shows distinct multi-voxel patterns for *clear* relative to the other operations, suggesting that this network diff in clearing information from WM. The Default Mode Network has distinct patterns for *suppress* and *clear*, also suggesting that clearing information from WM is distinct from suppressing it. The Frontoparietal Control Network displays distinct multi-voxel patterns for each of the four operations, suggesting that this network has high involvement in regulating the flow of information within WM. These results indicate that how information is removed from WM depends on distinct brain networks that each have a particular manner in which their co-activation patterns represent these operations.

**SIGNIFICANCE STATEMENT:** The ability to actively remove, manipulate and maintain information in working memory (WM) is required for encoding of new information and for controlling thoughts. This study revealed that different brain networks show characteristic multi-voxel activity patterns across four distinct WM operations: maintenance of information, replacement of one item by another, suppression of a specific item, and clearing the mind of all thought. One network, the Frontoparietal Control Network, differentiates all four operations, suggesting it may play a critical role in the controlled removal of information from WM.

## Introduction

The capacity of working memory (WM) is finite, and old information must be removed for new information to be effectively encoded (Ahmed & De Fockert, 2011; Lewis-Peacock et al., 2018). Most prior research has examined how information is gated into WM or how attention is shifted amongst items in WM, but not the mechanisms by which information is actively removed from WM. This question has been challenging to investigate because a method is needed to verify that an item in WM has actually been removed. Recent work using neuroimaging has overcome this problem and has provided insights into the mechanisms by which information can be removed and the brain regions that play a role in doing so (Banich et al., 2015; Kim et al., 2020).

This research has identified three distinct ways of removing information from WM: by replacing one item with another, by suppressing thoughts of a specific item, and by clearing the mind of all thoughts. Univariate analyses (Banich et al., 2015) revealed that some of these operations may involve the same brain regions. For example, compared to maintaining information all three of these operations activate superior parietal regions, consistent with the role of this region in shifting attention to information in WM (Tamber-Rosenau et al., 2011), while the prefrontal cortex is commonly activated for the suppress and clear operations, consistent with the idea that actively removing information is cognitively demanding.

Following up on these findings, Kim et al. (2020) used machine learning techniques to identify unique multivariate activation signatures for each of the four operations: *maintain, replace, suppress*, and *clear*. This approach reliably identifies regions that contribute to classifying the operations (i.e., classifier importance maps). Once again, it was observed that the same brain region contributes to multiple operations.

Given this overlap of activation across different operations in particular brain regions, it is not clear whether these regions are representing the operations in a unified manner (e.g., no distinction between *replace, suppress*, and clear) or whether these regions represent the operations in a distinct manner (e.g., distinguishing between each of *replace, suppress*, and *clear*). The current study examines this question. It does so from the perspective of identifying “representational brain networks,” which are a set of brain regions that show similar multi-voxel patterns of activation. This approach relies on representational similarity analysis (RSA), which identifies multi-voxel patterns of activity during a given mental operation (Kriegeskorte et al., 2008).

Here, we investigate whether there are distinct representational networks for WM removal operations. We re-analyzed the data from Kim et al. (2020) using RSA to evaluate the multi-voxel activation patterns within brain parcels (Glasser et al., 2016) for each of the four operations. Then Leiden community detection (Traag et al., 2019) was applied to these pattern matrices across the parcels to determine whether there are networks of brain regions that similarly represent those operations. Such an approach can provide important information. First, it can determine whether these WM operations are being differentially represented within brain regions, and second, it can identify whether there are certain brain networks that differentiate operations in a similar manner.

## Materials and Methods

Because this study involves a re-analysis of the data from Kim et al., (2020), full data collection procedures are described in that study and essential details are highlighted below.

### Participants

A total of 55 participants (17 male; age, M = 23.52, SD = 4.93) were included in all analyses. All participants had normal or corrected-to-normal vision and provided informed consent. The study was approved by the University of Colorado Boulder Institutional Review Board (IRB protocol # 16-0249).

### Stimuli

Stimuli for the fMRI study consisted of colored images (920 × 920 pixels) from three categories with three subcategories each: faces (actor, musician, politician), fruit (apple, grape, pear), and scenes (beach, bridge, mountain). Faces were recognizable celebrities, and scenes were recognizable locales (e.g., a tropical beach) or famous landmarks. Images were obtained from various resources, including the Bank of Standardized Stimuli and Google Images. Six images from each subcategory were used, for a total of 18 images per category and 54 images in total. All images were used for both the localizer and study phases of the experiment.

### Data Acquisition

MRI data were acquired on a Siemens PRISMA 3.0 Tesla scanner at the Intermountain Neuroimaging Consortium on the University of Colorado Boulder campus. Structural scans were acquired with a T1-weighted sequence, with the following parameters: repetition time (TR) = 2400 ms, echo time (TE) = 2.07 ms, field of view (FOV) = 256 mm, with a 0.8 × 0.8 × 0.8 mm^3^ voxel size, acquired across 224 coronal slices. Functional MRI (fMRI) scans for both the functional localizer and central study were obtained using a sequence with the following parameters: TR = 460 ms, TE = 27.2 ms, FOV = 248 mm, multiband acceleration factor = 8, with a 3 × 3 × 3 mm^3^ voxel size, acquired across 56 axial slices and aligned along the anterior commissure-posterior commissure line. For the functional localizer task, five runs were acquired in total, with each run consisting of 805 echo planar images (EPI), for a total of 4025 images across the five runs. For the central study, six runs were acquired in total, with each run consisting of a 1175 EPIs, for a total of 7050 images across the six runs. For full details of task and data acquisition, refer to Kim et al. (2020).

### fMRI Procedure

The experiment consisted of two phases completed in order: a functional localizer and a central study. Prior to completing both tasks in the MRI scanner, participants received training on the tasks outside of the scanner, including nine trials of the functional localizer task (three of each category of stimuli) and four self-paced trials of the central study (one trial per condition). Both tasks involved presenting participants with the same set of color images, though the tasks differed in what the participants were asked to do when presented with these images. All stimuli were presented on a black background with task-related words and fixation crosses shown in white font. Refer to Kim et al. (2020) for details on the five functional localizer runs. Note these runs were not included in the present analysis.

The central study was designed to allow us to track the representational status of a WM item while it was being manipulated using five distinct cognitive operations: maintaining an image in WM (maintain), replacing an image in WM with the memory of an image from a different subcategory of the same superordinate category (e.g., replacing an actor with a politician; replace subcategory), replacing an image in WM with the memory of an image from a different category (e.g., replacing an actor with an apple; replace category), suppressing an image in WM (suppress), and clearing the mind of all thought (clear). Note that classification results for replace subcategory and replace category trials from the previous study were nearly identical, and thus only replace category data are included in the main paper to control the number of trials across operation. On each trial, participants were presented with an image for 6 TRs (2760 ms) followed by another 6 TRs of an operation screen instructing participants how to manipulate the item in WM, and then a jittered inter-trial fixation lasting between 5 and 9 TRs (2300–4140 ms), consisting of a white fixation cross centered over a black background. The operation screen consisted of two words in the top and bottom halves of the screen, presented over a black background. For the maintain, suppress, and clear operations, the two words were the same: maintain, suppress, or clear, respectively. In the two replace conditions, the word in the top half was switch, whereas the word in the bottom half indicated the subcategory of image that the participant should switch to (e.g., apple). During practice and before the beginning of the task, participants were instructed to only switch to thinking about an image that had previously been shown during the functional localizer task. For example, if a participant was instructed to switch to thinking about an apple, that apple should be one of the apples that was presented to them during the localizer. Participants completed 6 runs of this task (9.01 min each, 54.05 min total), resulting in a total of 360 trials: 72 trials per operation, of which 24 trials were image category-specific trials per each operation condition. Each run had 40 TR long (18.4 s) fixation blocks at the beginning and end of the run. Within a single run, 12 trials were presented for each of the five operations, resulting in a total of 60 trials per run. Each image exemplar appeared at least once per operation condition across the entirety of the task. Trials were ordered pseudo-randomly within runs, with the order of trials optimized for BOLD deconvolution using optseq2 (Dale, 1999). We focused on four operations, and 288 trials were used in the main analysis.

### Analytic Overview

We identified communities (i.e., networks) of brain regions based on the RSA activation pattern similarities across four operations at the trial level to investigate the representational similarities and differences across operations regarding their underlying brain networks. The representational similarity between brain parcels (see below) allows us to investigate the brain’s community structure (networks) concerning the between-operations similarity in multi-voxel activation patterns. We chose RSA patterns to derive the brain networks because we aimed to investigate the similarity between brain regions regarding how they represent task-related information across the WM operations.

To derive our representational networks, we applied a multi-step network analysis procedure (Figure 1). First, the Glasser parcellation (Glasser et al., 2016) was used to extract 360 parcels covering a whole brain (180 parcels/hemisphere) for each subject. Next, activation patterns across all trials (i.e., 72 trial vectors for each of 4 operations yielding 288 trial vectors in total) were computed for each parcel. Trial vectors are weighted activation patterns that consist of multiple voxels (V_1_ ∼ V_N_: N = number of voxels in the parcel). The weighted activation value for each voxel equals the raw MR signal intensity multiplied by the GLM beta of the operation (operation > baseline) for the given voxel. The weighted measure of the pattern was used to ensure that only voxels that were sensitive to the WM operation were used and voxels that were primarily noise were removed.

**Figure 1.**
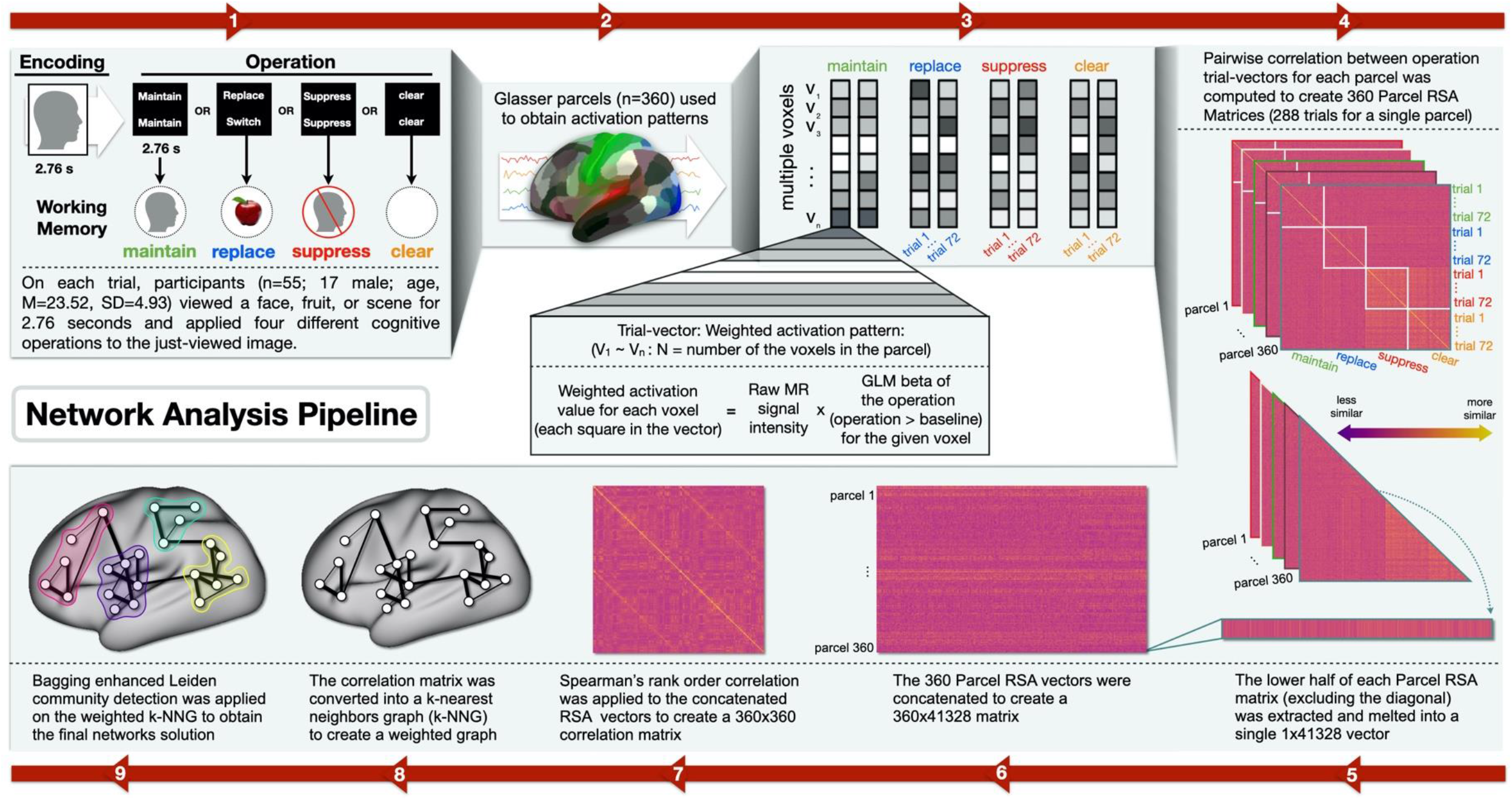
Outline of the steps to obtain the four representational brain networks. 1) On each trial, participants viewed a picture (a face, fruit, or scene) for 2.76 seconds (s). A screen appeared for 2.76 s indicating which one of four different cognitive operations (maintain, replace, suppress, clear) should be applied to the just-viewed image. 2) The Glasser parcellation (Glasser et al., 2016) was used to extract 360 parcels covering a whole brain (180 parcels/hemisphere) for each subject. 3) Activation patterns across all trials (i.e., 72 trial vectors for each operation and 288 trial vectors in total) were computed for each parcel. Trial vectors are weighted activation patterns that consist of multiple voxels (V_1_ ∼ V_N_: N = number of voxels in the parcel). The weighted activity value for each voxel equals the raw MR signal intensity multiplied by the GLM beta of the operation (operation > baseline) for the given voxel.4) Parcel RSA similarity matrices were then averaged across all subjects. 4) Pairwise correlation between operation trial vectors for each parcel was computed to create 360 Parcel RSA Matrices (288 trials for a single parcel). 5) The lower half of each Parcel RSA matrix was extracted and melted into a single 1×41328 vector. 6) These 360 Parcel RSA vectors are concatenated to create a 360×41328 matrix. 7) Spearman’s rank order correlation is then applied to the concatenated RSA matrices to create a 360×360 correlation matrix. 8) The 360×360 correlation matrix was converted into a weighted k-nearest neighbor graph (k-NNG) to generate the graph to derive our final representational networks. Weights for the k-NNG were obtained by calculating the Hadamard distances based on the overlap of neighbors (parcels) that represent the relationship between the pairwise similarity of the 360 parcels RSA patterns and their k-nearest neighbors. 9) Bagging enhanced Leiden community detection was applied on the weighted k-NNG to partition the graph to obtain our final (four) network solution.

Pairwise correlation between operation trial vectors for each parcel was computed to create 360 Parcel RSA Matrices (288 trials for a single parcel representing 72 trials per each of 4 operations), which provides information on the degree to which a given parcel is representing the operations in similar or dissimilar manner. We analyzed these data by adopting procedures that enable the construction of a weighted graph of similarity across the 360 brain parcels (Levine et al., 2015; Nikolaidis, Paksarian, et al., 2021; Shimoni, 2018). In this analysis, the lower half of each Parcel RSA matrix was extracted and melted into a single 1×41,328 vector (288 trials by 288 trials divided by 2; the diagonal and the top half is discarded as it is redundant with information in the lower half). These 360 parcel vectors were concatenated to create a 360×41,328 matrix. Spearman’s rank order correlations between each parcel vector was then computed to create a 360×360 correlation matrix. Finally, this correlation matrix was converted into a weighted k-nearest neighbor graph (k-NNG) that represented the similarity in activation patterns between parcels, where the weight of each edge represented the similarity between the connected parcels. Weights for the k-NNG were obtained by calculating the Hadamard distances based on the overlap of neighbors (parcels) that represent the relationship between the pairwise similarity of the 360 parcel vectors and their k-nearest neighbors. This approach circumvents arbitrarily thresholding the 360×360 correlation matrix to construct the distance metric before partitioning it to obtain the final networks. To prevent overfitting and bolster network robustness and reliability, k=36 was used, as determined using the “Sturges” formula, which implicitly bases k on the range of data (Nikolaidis, DeRosa, et al., 2021).

We chose a data-driven approach to derive our own networks based on patterns of activity across the four WM operations because pre-defined network templates may inaccurately reflect the organization of activity patterns in the brain across these operations. To determine whether the parcels could be grouped into networks that represent these operations in a similar manner, we applied Leiden community detection (LCD; Traag et al., 2019) on the weighted k-NNG to partition the graph to obtain our final networks solution. LCD has been used to derive well-connected structures in the brain (Hänisch et al., 2022) and has been proposed as a valid algorithm for deriving brain networks (Chen et al., 2022). To maximize network reproducibility, bagging (i.e., bootstrap aggregation) was used in our LCD analysis (Nikolaidis et al., 2020). We then performed additional analyses with regard to the similarity and/or dissimilarity of activation patterns within each of the four representational networks identified by LCD. More details on each of these steps are provided below.

### Representational Similarity Analysis

The Glasser parcellation was converted from the MNI space to native space for each subject, and the multivoxel activation patterns were extracted for each parcel in the native space. For each trial, the activation patterns were averaged across the operation period (i.e., 2.75 s, 6 TRs) from the onset that was shifted forward 4.6 s (10 TRs) to account for hemodynamic lag. To emphasize the important voxels, the averaged patterns were weighted with the beta estimate contrast from the general linear model (GLM) analysis. In the GLM model, four operations, utilizing a canonical hemodynamic response function, and six motion regressors were modeled. Each operation weight was obtained from the t-contrast (e.g., maintain > others) without thresholding. The similarity of the weighted patterns across trials (i.e., 288 trial vectors total, 72 trials/operation) were then computed with Pearson’s correlation (i.e., RSA) to create the operation similarity matrix (288 × 288) for each parcel. Fisher’s z transformation was applied on the correlation coefficient to average the patterns across subjects into a group similarity matrix. The bottom of the off-diagonal operation similarity matrix for each parcel, which indicates the similarity and dissimilarity across four operations in the trial level for each parcel, was used for the LCD analysis.

### Leiden Community Detection for deriving representational networks

LCD is a data-driven clustering method applied in this study to identify groups of brain regions with similar activation patterns across working memory (WM) operations. This algorithm is well-suited for this task due to its three-phase community detection procedure, which is based on hierarchical clustering. The algorithm aims to maximize the modularity (Q) of the communities, which is a quantitative measure of the number of edges found within communities compared to those predicted in a random graph with an equivalent degree distribution. A positive Q value indicates a reliable division of communities into disjoint groups, with the highest Q value representing the optimal community structure.

The LCD algorithm starts with a single graph partition and iteratively refines it by moving individual parcels from one community to another until no further improvements can be made. This process is repeated until the optimal community structure is reached. An attractive feature of the LCD algorithm is its ability to determine the optimal number of clusters, which can help prevent incorrect cluster approximation that may occur with other methods, such as hierarchical or k-means clustering. Furthermore, the algorithm uses the Q function as its objective function, which aims to maximize intra-community similarity while minimizing inter-community similarity.

This feature is particularly useful in the present analysis as it allows for a data-driven determination of the most appropriate number of clusters and helps ensure that the resulting clusters accurately reflect the underlying structure of the data. Overall, the LCD algorithm provides a robust and reliable method for identifying groups of brain regions with similar activation patterns, making it an ideal choice for this type of analysis.

Initially, we conducted two separate LCD analyses that used Spearman rank-order correlation (SRC) and Euclidean distance for the distance measure for k-NN. SRC was chosen as the more appropriate distance metric for the k-NNG graph because the LCD outputs when SRC was used yielded higher modularity (Q), indicating higher community clustering (SRC BaggedQ=.653; Euclidean BaggedQ=.447) and stronger parcel allegiance/reproducibility (SRC meanARI=.88 **±** .07; Euclidean meanARI=.67 **±** .13) indicating stable community assignment. Q was also examined across each of the 1000 bootstrapped iterations, and it was revealed that, on average, SRC had higher Q per iteration (meanQ=.53 **±** .02) compared to Euclidean (meanQ=.21 **±** .02).

### Bootstrap aggregation (Bagging) for representational network robustness and reliability

To identify and verify the most stable communities, a bagging approach was used. Bagging begins by resampling the concatenated 360×41,328 parcel vectors matrix with replacement (i.e., bootstrapping) and aggregating across bootstrap samples. This technique reduces variability in the estimation process by averaging across multiple resampled datasets. When applied to clustering, bagging has improved cluster robustness and reliability (Nikolaidis et al., 2020). The fundamental advantage of bagging stems from the additional value of combining multiple cluster assignments into a single clustering solution. More specifically, the features for ensemble clustering consist of the aggregation of cluster outputs themselves. LCD is then applied, resulting in cluster solutions for every bootstrap iteration. Each cluster solution is transformed into individual adjacency matrices that are summed together to create an adjacency matrix (similarity matrix) of the total number of times participants were in the same cluster. Next, mask adjacency matrices are simultaneously created and summed to equate the total number of times participants went into the same LCD iteration together (inclusion matrix). The similarity matrix is then divided by the inclusion matrix to create a mean adjacency matrix (stability matrix) that is then turned into a weighted network. Finally, LCD is applied one more time to obtain the final cluster solution.

### Multidimensional scaling for representational dissimilarity

To investigate the similarities of activation patterns across representational networks, we applied multidimensional scaling (MDS) on the operation dissimilarity matrix to compute the distance across parcels and across the four operations in each community (Derndorfer & Baierl, 2013). This provides a visual representation of a complex set of relationships between activation patterns that can be visually interpreted to understand the pattern of proximities (i.e., similarities or distances) among the parcels and operations. MDS is a mathematical operation that converts an item-by-item matrix (e.g., 288 × 288 operation dissimilarity matrix) into an item-by-variable matrix (e.g., 288 × 3 matrix). The input to MDS is a square, symmetric 1-mode RSA pairwise dissimilarity matrix that contains correlations across the connectivity patterns from continuous time points. The dissimilarity was transformed by subtracting the pairwise correlation coefficient value from one and then dividing the difference by two, resulting in a distance value that ranges from zero to one. Each observation is then assigned coordinates in each of the dimensions. Three dimensions were chosen to maximize the interpretation of the high-dimensional RSA connectivity patterns.

There are three important considerations for our MDS representational space. The first is that the axes (dimensions) are meaningless and thus the orientation within the space is arbitrary. Therefore, our MDS representation of distances between the RSA connectivity patterns by parcel need not be oriented such that positive values indicate more activation and vice versa. Second, the purpose of the MDS space is to identify which points are close to others, and we interpret points that are close to one another as having similar connectivity patterns. Third, the three substantive dimensions need to be different from the mathematical dimensions (axes) that define the MDS vector space. For example, the three specified dimensions used to generate similarities may be much larger than the number of dimensions needed to reproduce the observed pattern. This is because the three dimensions are purposefully orthogonal, making them maximally efficient.

### Uniqueness of WM operation patterns

To evaluate the accuracy of classifying the MDS operation similarity matrix patterns for each representational network, we performed bootstrapped multiclass classification using a support vector classifier (SVC) from Scikit-learn in Python (Pedregosa et al., 2012). The patterns were analyzed using an across-operation classification and a pairwise-operation classification approach. Ten K-fold cross-validations indicated SVC had the highest performance with a radial basis function kernel, gamma parameter set to ‘scale’ and the regularization parameter (C) set to 1. All subsequent classification models used these parameters.

Across-operation classification evaluated our multiclass models by comparing each operation (class) against all the others simultaneously. For each classifier iteration, 288 bootstrap-resampled MDS connectivity patterns were fed into the classifier. Using a 1×4 pattern design, we take a given WM operation and consider it as the “positive” class, while the other three WM operations are considered as the “negative” class. A 1×4 pattern design in our across-operation (also known as one vs. rest) classification refers to a strategy for separating multi-class problems into multiple binary classification problems. In the 1×4 pattern, SVM separates the data into four binary classification problems, where each problem compares one of the four WM operations and all the other three WM operations combined into a single group. This procedure uses all the known binary classification metrics to evaluate this scenario.

The pairwise-operation classification compares all possible pairwise combinations of WM operations (e.g., Suppress vs Clear, Suppress vs. Maintain). The first step creates a dataset containing only the two operations being examined and consists of 144 bootstrap-resampled MDS connectivity patterns. Then we define observations with real class = “WM operation 1” as our positive class and the ones with real class = “WM operation 2” as our negative class. Note that “WM operation 1 vs WM operation 2” is different than “WM operation 2 vs WM operation 1”, so both cases were accounted. Because of this, we produced 12 unique pairwise-operation classification scores from all pairwise combinations of the four operations.

The Precision and Recall (PR) Curve and Area Under the Curve (AUC) were chosen as the metrics for our multiclass classification. The PR curve allowed us to assess the relationship between the precision (the number of true positives over the total number of positives - true positives and false positives combined) and the recall (number of true positives over the total number of real true values - the true positives and false negatives combined). Specifically, precision assesses the number of times our model correctly classified each WM operation’s connectivity pattern to all of the positive classifications by our model. Recall assesses the ratio of correct positively classified operation connectivity patterns to the total number of connectivity patterns that should have been predicted as positive. Higher recall indicates that more correctly positive operation connectivity patterns were detected. The PR Curve and the AUC score are important tools for evaluating binary classification models. They reveal the separability of the WM operations by evaluating how well the SVC model classified each operation.

## Results

### Representational Network Identification

We derived the primary representational network communities underlying the implementation of the WM operations for *maintain, replace, suppress*, and *clear*. LCD uncovered four stable networks (Figure 1E), similarly representative of previously established functional networks (Greene et al., 2016; Hearne et al., 2016; Yeo et al., 2011). The labels for our networks were derived based on their highest correlations with the labels of the Yeo 7 networks (Yeo et al., 2011). Correlations were established between dummy-coded voxels corresponding to each of our four and Yeo 7 networks. The highest correlations for one of our networks spanned the Frontoparietal, Ventral Attention, and Dorsal Attention from the Yeo7 and thus was dubbed the Frontoparietal Control Network (FPCN). A list of the Glasser parcels comprising each of our derived networks can be found <u>here</u>.

#### Cluster 1 – Visual Network (VN)

consists of 76 parcels that are mostly located within the Visual cortex (Primary, Dorsal Stream, Early, Complex, and Neighboring) and Superior Parietal lobe and, for purposes of the present analysis, was labeled as the Visual Network.

#### Cluster 2 – Somatomotor Network (SMN)

consists of 63 parcels primarily located in the Somatosensory cortex, Motor cortex, Early Auditory cortex, Paracentral Lobular cortex, and Mid-Cingulate cortex.

#### Cluster 3 – Default Mode Network (DMN)

consists of 121 parcels primarily located in the Posterior Cingulate, Anterior Cingulate, Medial Prefrontal cortex, Insular cortex, Frontal Opercular cortex, Orbital cortex, Polar Frontal cortex, Inferior cortex, and Superior Parietal cortex.

#### Cluster 4 – Frontoparietal Control Network (FPCN)

consists of 100 parcels primarily located in the Dorsolateral Prefrontal cortex, Inferior Frontal cortex, Superior Parietal cortex, Inferior Parietal cortex, Anterior Cingulate cortex, and Medial Prefrontal cortex.

### Multidimensional Representational Network Patterns

We compared the multi-voxel activation patterns across and within networks for each of the four WM operations to determine how each operation is represented. To begin, we first averaged the RSA operation similarity matrices for the parcels belonging to each network. This analysis resulted in four unique network RSA operation similarity matrices (**Figure 2B**). These matrices were then concatenated together and reduced to three dimensions via MDS for visualization (**Figure 3A**). Across-operation classification revealed that each network’s similarity patterns (across operations) were highly dissociable from each other’s (AUC range 0.94-1.0). We then repeated this MDS analysis on each network’s averaged RSA operation similarity matrix to evaluate the uniqueness of each WM operation’s activation pattern (**Figure 3B-E**).

**Figure 2.**
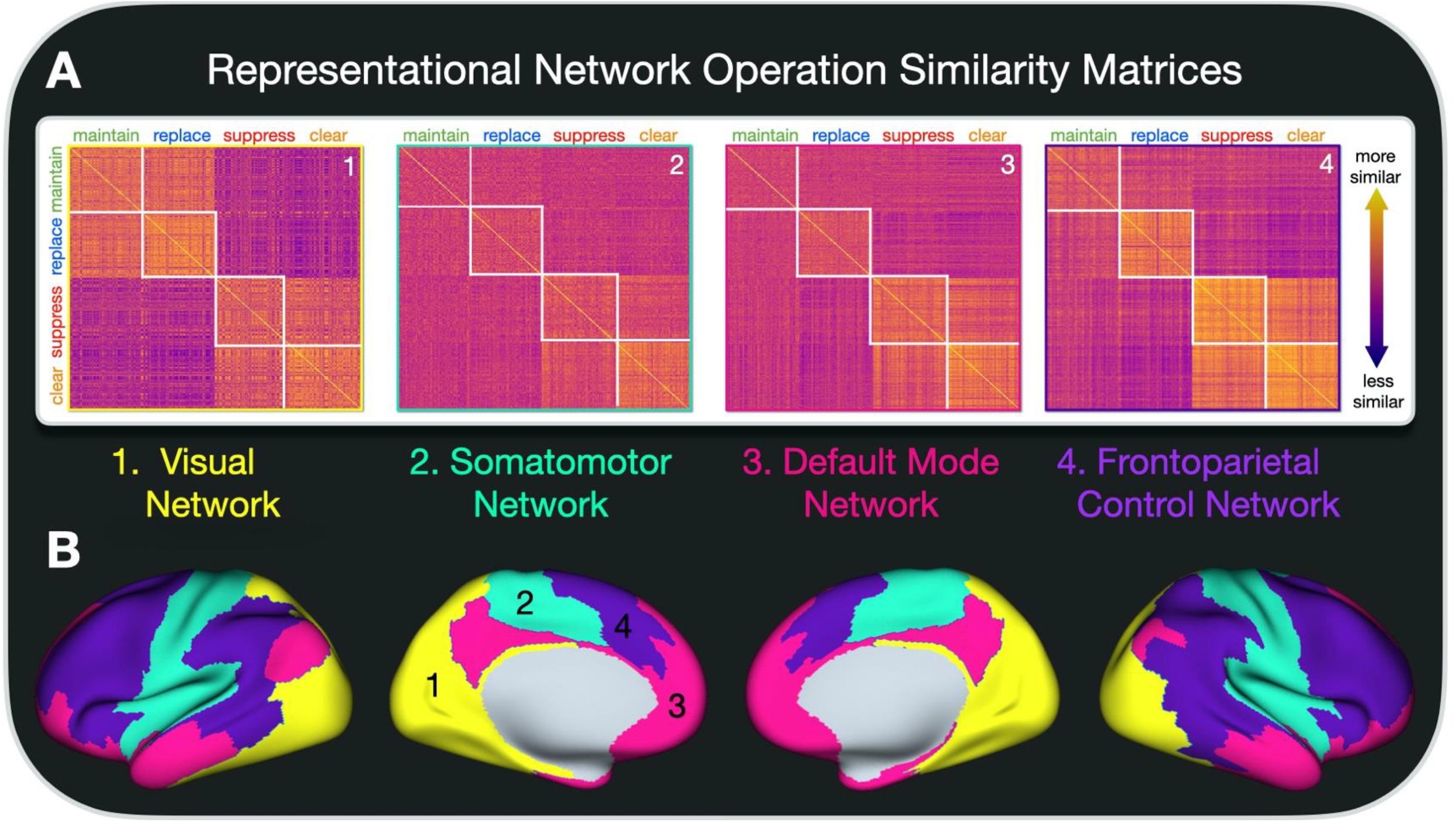
Representational network profiles. A) Representational network RSA operation similarity matrices correspond to the individual parcel RSA operation similarity matrices for each community (network) averaged together. Numbers and outline color correspond to each network. B) Glasser parcels are colored by network allegiance – 1. Visual Network, yellow; 2. Somatomotor 3. Network, teal; 4. Default Mode Network, pink; Frontoparietal Control Network, purple.

**Figure 3.**
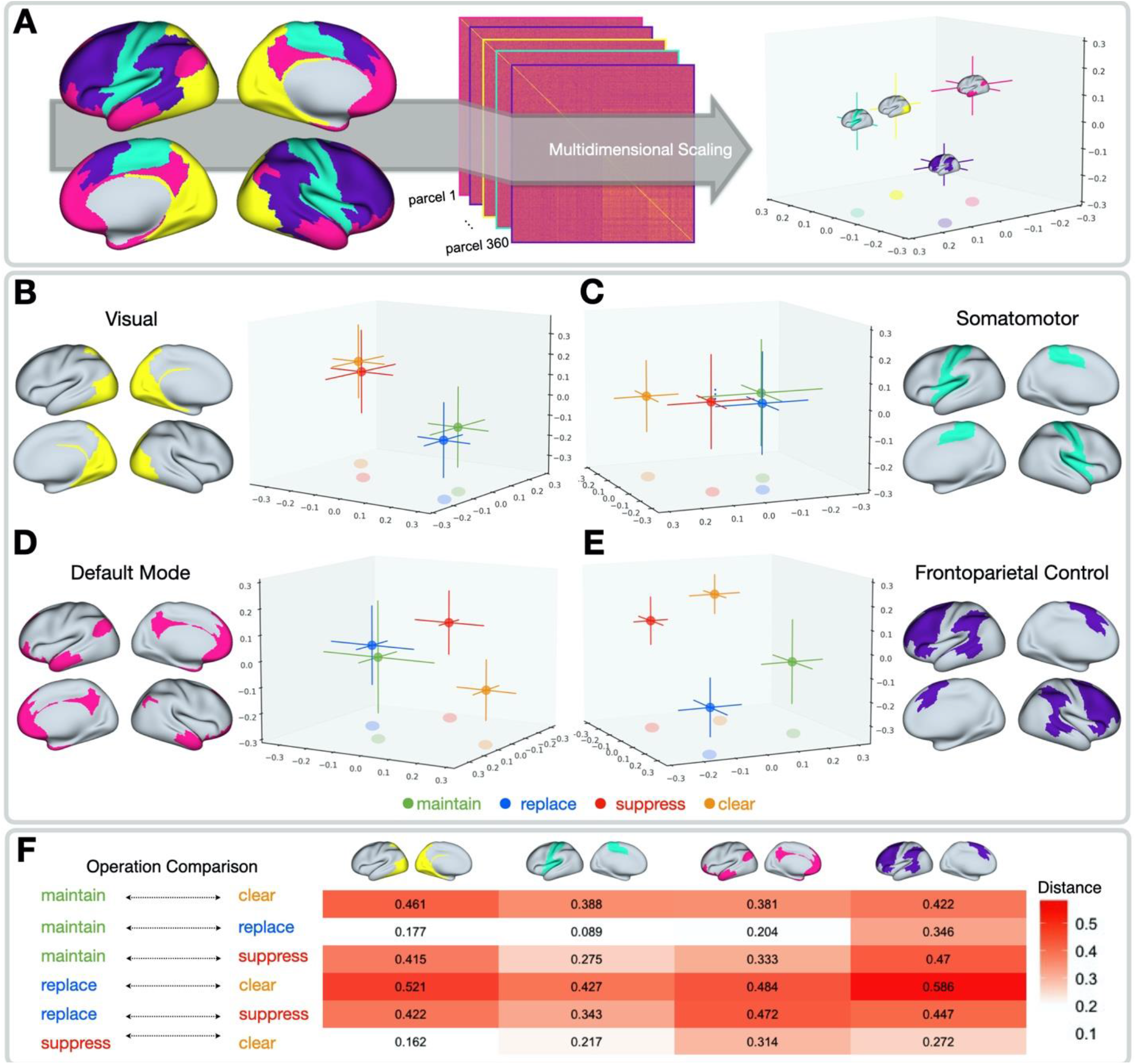
Multidimensional scaling of the activity patterns for each of the four WM operations between and within network. A) Brains are colored by network (Visual Network, yellow; Somatomotor Network, teal; Default Mode Network, pink; Frontoparietal Control Network, purple) and represent the mean, and bars represent the mean standard error of the 95% confidence interval across each parcel’s representational patterns for all of the operations. B-E) Within network operation MDS profiles, B) Visual Network, C) Somatomotor Network, D) Frontoparietal Network, E) Default Mode Network. Single points represent the mean, and bars represent the mean standard error of the 95% confidence interval across the MDS representational patterns between trial vectors across the 288 trials (72 for each operation) for each parcel within a given network and colored by operation (maintain, green; replace, blue; suppress, red; clear, orange). F) Euclidean distances between the pairwise Mean MDS pattern across each operation by network.

Next, across-operation classification and pairwise-operation classification were computed for each network’s multi-voxel representational patterns for each WM operation (Table 1). Below we review each network’s representational patterns and the across-operation classification and pairwise-operation classification analysis results. Finally, the pairwise distance was calculated between the mean operation representational patterns across the three MDS coordinates (**Figure 3F**). Higher distances indicate greater separation between the operations and thus indicate that the network elicits different activity patterns for performing each operation. Conversely, smaller distances indicate that the network does not elicit unique activity patterns to perform each operation, implying that the network does represent those operations in distinct manners.

**Table 1.**
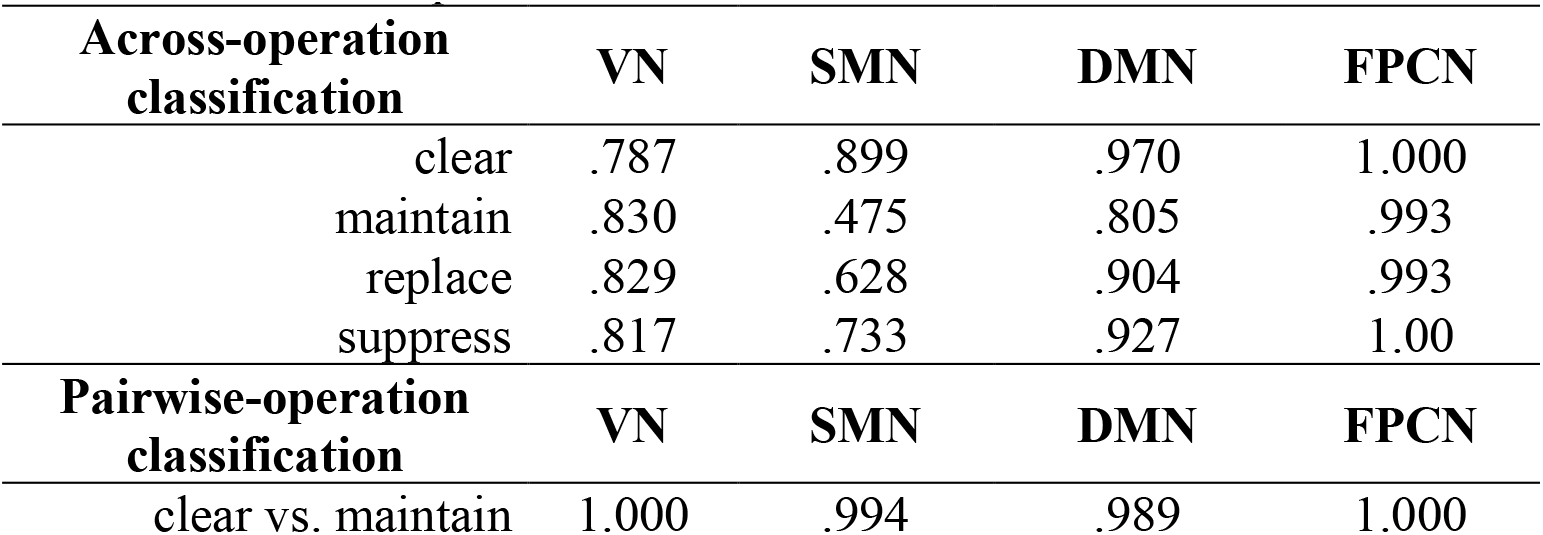
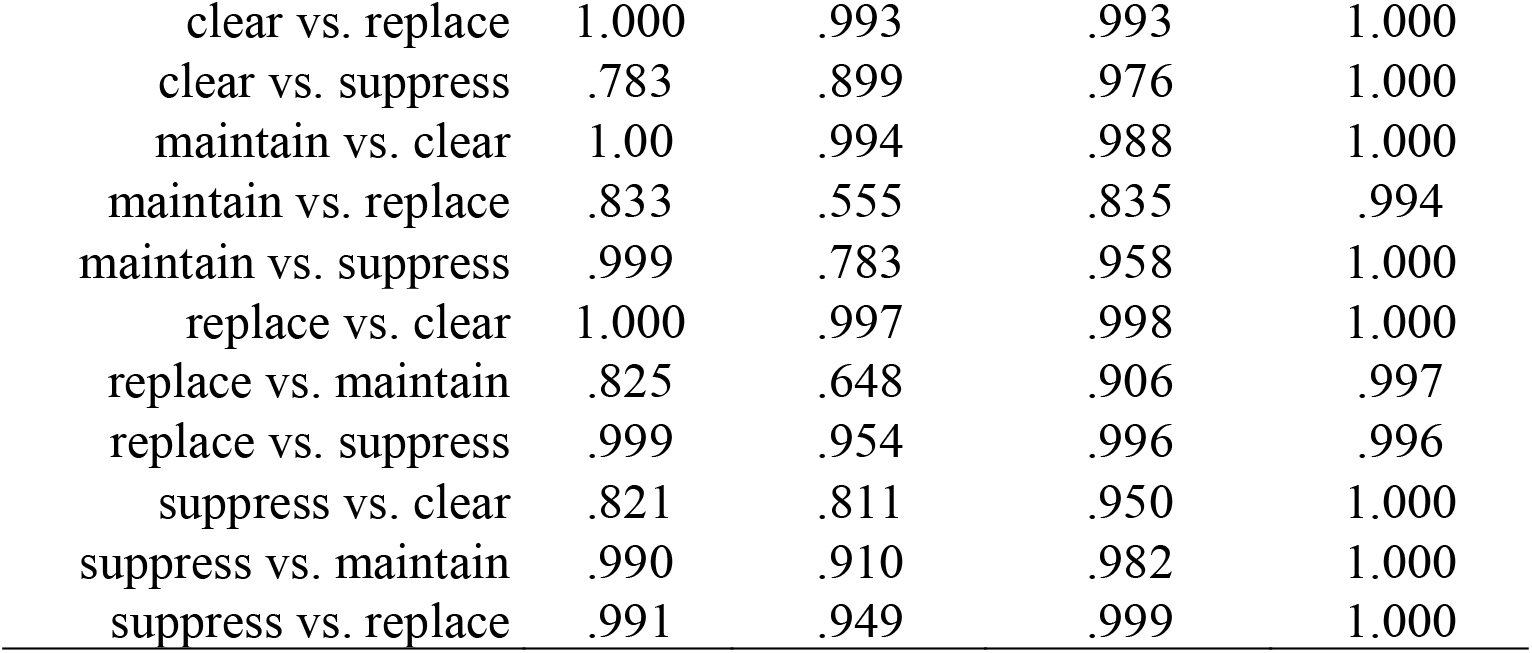
Network Operation Precision and Recall Area Under the Curve

| **Across-operation**<br>**classification** | **VN** | **SMN** | **DMN** | **FPCN** |
| --- | --- | --- | --- | --- |
| clear | .787 | .899 | .970 | 1.000 |
| maintain | .830 | .475 | .805 | .993 |
| replace | .829 | .628 | .904 | .993 |
| suppress | .817 | .733 | .927 | 1.00 |
| **Pairwise-operation**<br>**classification** | **VN** | **SMN** | **DMN** | **FPCN** |
| clear vs. maintain | 1.000 | .994 | .989 | 1.000 |
| clear vs. replace | 1.000 | .993 | .993 | 1.000 |
| clear vs. suppress | .783 | .899 | .976 | 1.000 |
| maintain vs. clear | 1.00 | .994 | .988 | 1.000 |
| maintain vs. replace | .833 | .555 | .835 | .994 |
| maintain vs. suppress | .999 | .783 | .958 | 1.000 |
| replace vs. clear | 1.000 | .997 | .998 | 1.000 |
| replace vs. maintain | .825 | .648 | .906 | .997 |
| replace vs. suppress | .999 | .954 | .996 | .996 |
| suppress vs. clear | .821 | .811 | .950 | 1.000 |
| suppress vs. maintain | .990 | .910 | .982 | 1.000 |
| suppress vs. replace | .991 | .949 | .999 | 1.000 |

**Table 1.** Network Operation Precision and Recall Area Under the Curve
| **Across-operation**<br>**classification** | **VN** | **SMN** | **DMN** | **FPCN** |
| --- | --- | --- | --- | --- |
| clear | .787 | .899 | .970 | 1.000 |
| maintain | .830 | .475 | .805 | .993 |
| replace | .829 | .628 | .904 | .993 |
| suppress | .817 | .733 | .927 | 1.00 |
| **Pairwise-operation**<br>**classification** | **VN** | **SMN** | **DMN** | **FPCN** |
| clear vs. maintain | 1.000 | .994 | .989 | 1.000 |
| clear vs. replace | 1.000 | .993 | .993 | 1.000 |
| clear vs. suppress | .783 | .899 | .976 | 1.000 |
| maintain vs. clear | 1.00 | .994 | .988 | 1.000 |
| maintain vs. replace | .833 | .555 | .835 | .994 |
| maintain vs. suppress | .999 | .783 | .958 | 1.000 |
| replace vs. clear | 1.000 | .997 | .998 | 1.000 |
| replace vs. maintain | .825 | .648 | .906 | .997 |
| replace vs. suppress | .999 | .954 | .996 | .996 |
| suppress vs. clear | .821 | .811 | .950 | 1.000 |
| suppress vs. maintain | .990 | .910 | .982 | 1.000 |
| suppress vs. replace | .991 | .949 | .999 | 1.000 |

### Visual Network

The visual network revealed two different activity patterns across the four operations. *Maintain* and *replace* showed similar patterns across trials but varied considerably from *suppress* and *clear*, which elicited similar patterns (**Figure 2B** network community 1; **Figure 3B**). Across-operation classification for each operation was moderate (0.787-0.83). However, pairwise-operation classification revealed higher classification across *maintain*/*replace* to *suppress*/*clear* (0.99-1.00) compared to between them (.783-.833). The distance between the operation activity patterns and the pairwise-operation classification AUCs were highly correlated across the operation comparisons (r=.967, p < .001). See Table 1 for all across-operation and pairwise-operation classification PR Curve AUC scores and Figure 3F for all mean MDS distances between the operations by network.

### Somatomotor Network

The SMN revealed non-distinct activity patterns across the operations, except for *clear* (**Figure 2B** network community 2; **Figure 3C**), and to a lesser degree *suppress*. R*eplace, maintain*, and *suppress* revealed the lowest across-operation classification AUCs (.475-.678) compared to *clear* (.899). Notably, the pairwise-operation classification AUCs were the lowest between *maintain* and *replace* (.555-.648) as compared to *suppress* (.*783-*.*954*) and *clear* (.*899-*.*994)*. The mean pairwise distance between the operation activity patterns and the pairwise-operation classification AUCs were highly correlated across the operation comparisons (r=.931, p < .001).

### Default Mode Network

The DMN revealed two unique activity patterns for each of the *clear* and *suppress* operations, as shown in **Figure 2B** (network community 3) and **Figure 3D** (Across-operation classification AUCs: 0.927-0.97; and pairwise-operation classification AUCs: 0.95-.976). The distance between the operation’s activity patterns and the pairwise-operation classification AUCs was also reasonably correlated across the operation comparisons (r=.841 p = .001).

### Frontoparietal Control Network

The FPCN revealed dissociable activity patterns for each of the four operations and showed the highest across-operation classification AUCs (0.993-1.00) and pairwise-operation classification AUCs (0.994-1.00) amongst the four networks of interest, as one might expect for a network known for exerting control (**Figure 2B** network community 4; **Figure 3E**). The correlation between the pairwise operation distance and the pairwise operation AUCs was negligible (r=0.261, p=.41). The operation AUCs for the FPCN were all > .99. Despite the operation distances being predominantly larger on average than those of other networks, the low variability for each of the operations was the primary factor contributing to the performance of the classifier scores, rather than the distances, which varied to some extent across operations in the FPCN.

## Discussion

The present study revealed that four major brain networks contribute to removing information from WM. Based on the similarity of the multi-voxel activity patterns, we identified that the Visual Network, Somatomotor Network, Default Mode Network, and Frontoparietal Control Network are uniquely involved in removal. Next, we discuss the patterns for each network and what these patterns reveal about control over the contents of working memory.

### Visual Network (VN)

Within the VN, there was a clear distinction between the activity patterns observed for the *maintain* and *replace* operations, which require information to be held in WM, compared to the *suppress* and *clear* operations, which require the currently held information to be removed entirely. These findings are highly consistent with the univariate results from Banich et al. (2015) that showed evidence of activity in the ventral visual processing stream for the *maintain* and *replace* operations but no significant activity above a fixation baseline for the *suppress* and *clear* operations. The current results expand upon those findings to demonstrate that distinct activity patterns separate the *maintain* and *replace* operations from those during the *suppress* and *clear* operations.

### Somatomotor Network (SMN)

The SMN network revealed a representational pattern for clearing the mind of all thoughts that was highly distinguishable from the other operations. To a lesser degree, *suppres*s also reveal patterns separable from *maintain* and *replace*. We can only speculate why *clear* and *suppress* produced this pattern. One suggestion offered by Banich et al. (2015) about the unique univariate brain activation pattern observed for *clear* is that clearing the mind completely of all thoughts results in a brief attentional shift away from external sensory processing to a focus on one’s internal states. Our findings align with this idea. Additionally, previous work using transcranial magnetic stimulation suggested that the SMN can directly inhibit neural activity in regions involved in maintaining or representing information to be suppressed (Romei et al., 2010). In the *maintain* and *replace conditions*, there is a higher amount of sensory representation that must be held or manipulated, which makes the lack of differences across these operations unsurprising, considering the SMN has been primarily implicated in processing sensory information (Uddin et al., 2019). The SMN may also interact with other networks, such as the FPCN and DMN, and further, modulate neural activity in regions that remove information in WM. Further research is needed to understand the mechanisms underlying the processes within the SMN fully.

### Default Mode Network (DMN)

Like the SMN, the DMN exhibits a mixture of representational patterns that distinguish between the *clear* and *suppress* operations, compared to those of *maintain* and *replace* operations. However, unique to the DMN, *suppress* exhibits a more sharply delineated pattern from *maintain* and *replace*. Like the VN, the DMN distinguishes operations that involve holding information in WM, *maintain* and *replace*, from those which require information to be removed entirely, *suppress* and *clear*. This distinction is consistent with findings that activity of the DMN is observed more often when cognitive processes are more inwardly directed as compared to oriented toward external stimuli (Andrews-Hanna et al., 2014; Kucyi et al., 2016; Raichle, 2015). Unlike the VN, however, there is also a distinction between the *suppress* and *clear* operations (consistent with Banich et al. (2015) and Kim et al., (2020), which suggests that the DMN may have an integral role in distinguishing how information is removed from mind.

### Frontoparietal Control Network (FPCN)

The FPCN is the only network that showed four distinct representational patterns for each operation, suggesting this network may be important for removing information from WM. This notion is consistent with previous work demonstrating the FPCN’s involvement in WM tasks (Finc et al., 2020). Our findings are consistent with the role of the FPCN in various executive control operations (Marek & Dosenbach, 2018; Scolari et al., 2015), and with the idea that the FPCN uses representational codes to flexibly distinguish between each operation much as they do to distinguish between types of information in WM (Nee & Brown, 2012).

One aspect of these results to consider, given that representations of these operations seem to cohere across different brain networks, is the degree to which such representational similarity is supported by connectivity and communication within these brain regions. While the current study cannot address these issues, such coherence likely occurs via connectivity as recent research has found significant overlap between brain regions that represent information in similar manners (representational network) and brain regions that form networks based on their pattern of connectivity (Pillet et al., 2019).

The present work also adds to our knowledge of the neural underpinnings with regards to prior work by our group. Banich et al. (2015) employed mass-univariate analyses of fMRI data to show that a hierarchy of brain regions implicated in cognitive control is engaged depending on the operations used to remove information from WM. Kim et al. (2020) extended these findings with multivariate analyses applied to brain activation patterns that 1) verified that specific information was removed from WM, 2) identified the consequences of each operation on the representation of the information being removed, and 3) assessed the impact of the operation on the encoding of new information in WM. The current findings build off these studies by showing that brain regions do not work in isolation to remove information from WM. Instead, there are distinct networks that each represent these operations similarly but in ways distinct from other networks.

### Limitations and Future Directions

While the current results identified four representational networks, each of which revealed insight into their role in regard to the WM operations investigated, this study has its limitations. This study is limited by its ability to obtain a more mechanistic understanding of how the representational patterns within each network implement the operations. For example, they do not determine the degree to which each of these representational patterns within a network are causally related to the execution of each operation. One possibility is that the operation is actually implemented based on the combined representational pattern across each of the four networks (e.g., *maintain* = representational pattern for *maintain* for the SMN + VN + DMN + FPCN). An alternative possibility is that the operations are implemented primarily by the FPCN, as it is the only network that distinguishes between the four operations, and the representational patterns observed in the other networks are a by-product of top-down control. One way to address this issue is to determine the degree to which the representational pattern of a given network or combination of networks can predict the degree to which the representation of given item is effectively removed from WM, as potentially indexed by classifier fits or RSA of specific items, as identified in Kim et al., (2020).

One future direction is to investigate the degree to which the representational patterns of the operations are robust at the level of an individuals, and whether such differences may be linked to behavioral characteristics. Individual differences in brain network topography and topology have been compared to group-average estimates (Gordon et al., 2017), and those individual differences in network topography have been linked to individual differences in levels of executive functioning (Cui et al., 2020). Whether there are stable individual differences in the representation of these operations as measured by the metrics employed here and whether they can be linked to behavioral measures of removing information from WM and/or self-report of difficulties in controlling thoughts remains to be seen. It may be that individuals whose representations of the operations in the FPCN are not clearly distinguished might have difficulty in removing information in WM. Such examinations are of interest as characteristics of activation and connectivity of DMN and FPCN have been linked with many forms of psychopathology (Bullmore & Sporns, 2009; Garrity et al., 2007; Harrison et al., 2007) and internalizing symptomology (Banich et al., 2020). If representational patterns within these networks were found to be associated with someone’s inability to effectively control the contents of WM, future investigations might use these patterns to assess the effectiveness of interventions and/or for use in cognitive training and biofeedback to normalize the representational patterns between the regions of these network (Pandria et al., 2018; Van den Boom et al., 2018).

### Conclusion

This study leveraged various computational approaches to identify networks of brain regions whose activity differentiates when someone *maintains, replaces, suppresses*, or entirely *clears* information from mind. Four networks were identified that have distinct configurations across the four WM operations based on the similarity of their activation patterns. Notably, the FPCN revealed distinct representational patterns for each operation, suggesting the regions within this network differentiate these four operations. Additionally, the DMN revealed two distinct patterns for *suppress* and *clear*, suggesting that the regions within this network distinguish between these two methods of removing information held in WM. These findings provide new insights into the neural underpinnings of information removal from WM and pave the way for future studies to explore the stability of representational patterns across individuals and their potential application for cognitive training and intervention.

